# A single mechanism for global and selective response inhibition under the influence of motor preparation

**DOI:** 10.1101/2020.03.16.993261

**Authors:** Liisa Raud, René J. Huster, Richard B. Ivry, Ludovica Labruna, Mari S. Messel, Ian Greenhouse

## Abstract

In our everyday behavior, we frequently cancel one movement while continuing others. Two competing models have been suggested for the cancellation of such specific actions: 1) the abrupt engagement of a unitary global inhibitory mechanism followed by reinitiation of the continuing actions, or 2) a balance between distinct global and selective inhibitory mechanisms. To evaluate these models, we examined behavioral and physiological markers of proactive control, motor preparation, and response inhibition using a combination of behavioral task performance measures, electromyography, electroencephalography, and motor evoked potentials elicited with transcranial magnetic stimulation. Healthy participants performed two versions of a stop signal task with cues incorporating proactive control: A unimanual task involving the initiation and inhibition of a single response, and a bimanual task involving the selective stopping of one of two prepared responses. Stopping latencies, motor evoked potentials, and frontal beta power (13-20 Hz) did not differ between the uni- and bimanual tasks. However, evidence for selective proactive control before stopping was manifest in the bimanual condition as changes in corticomotor excitability, mu (9-14 Hz), and beta (15-25 Hz) oscillations over sensorimotor cortex. Altogether, our results favor the recruitment of a single inhibitory stopping mechanism with the net behavioral output depending on the levels of action-specific motor preparation.

**Significance statement:** Response inhibition is a core function of cognitive flexibility and movement control. Previous research has suggested separate mechanisms for selective and global inhibition, yet the evidence is inconclusive. Another line of research has examined the influence of preparation for action stopping, or what is called proactive control, on stopping performance, yet the neural mechanisms underlying this interaction are unknown. We combined transcranial magnetic stimulation, electroencephalography, electromyography and behavioral measures to compare selective and global inhibition models and to investigate markers of proactive control. The results favor a single inhibitory mechanism over separate selective and global mechanisms, but indicate a vital role for preceding motor activity in determining whether and which actions will be stopped.

## Introduction

Flexible movement control requires the ability to inhibit a specific component of multiple actions. Response inhibition can be studied with the stop signal task, where the primary instruction is to respond as quickly as possible to a go signal. On a fraction of trials, a stop signal is presented after the go signal and participants are instructed to inhibit their response. In a bimanual-selective version of this task, the go signal requires simultaneous responses with both hands and the stop signal requires inhibition of only one hand’s response.

A way to dissociate inhibition mechanisms in the uni- and bimanual tasks is to compare putative inhibition indices. A behavioral index of stopping latency is the stop signal reaction time (SSRT), estimated based on the independent horse race model (Logan and Cowan, 1984; Band et al., 2003). A peripheral measure of inhibition latency is the partial response electromyography (prEMG), detectable when an initiated response is interrupted (Raud and Huster, 2017). Motor evoked potentials (MEPs), elicited with transcranial magnetic stimulation (TMS) over the motor cortex, provide a physiological index of corticospinal excitability during response preparation, execution, and inhibition (Bestmann and Krakauer, 2015). Electroencephalography (EEG) provides further measures of cortical activity, with the stop signal evoking power increase in the beta band (13-20 Hz) over frontal electrodes (Picazio et al., 2014; Wagner et al., 2018).

Two models have been offered to explain selective stopping (Figure1). One proposes that the motor system is globally inhibited via the hyperdirect cortical-subthalamic nucleus pathway (Wessel and Aron, 2017). Reduced MEP amplitudes in task-irrelevant muscles during stopping support such global inhibition (Badry et al., 2009; Cai et al., 2012; Greenhouse et al., 2012; Majid et al., 2012; Wessel et al., 2013). In the bimanual task, selective stopping can be achieved by the global inhibitory signal to interrupt all movements followed by re-activation of the required response (Coxon et al., 2007; Macdonald et al., 2014; Cowie et al., 2016). This inflates reaction times (RT) of the responding hand and causes a transient change in its EMG profile (Macdonald et al., 2012).

**Figure 1.**
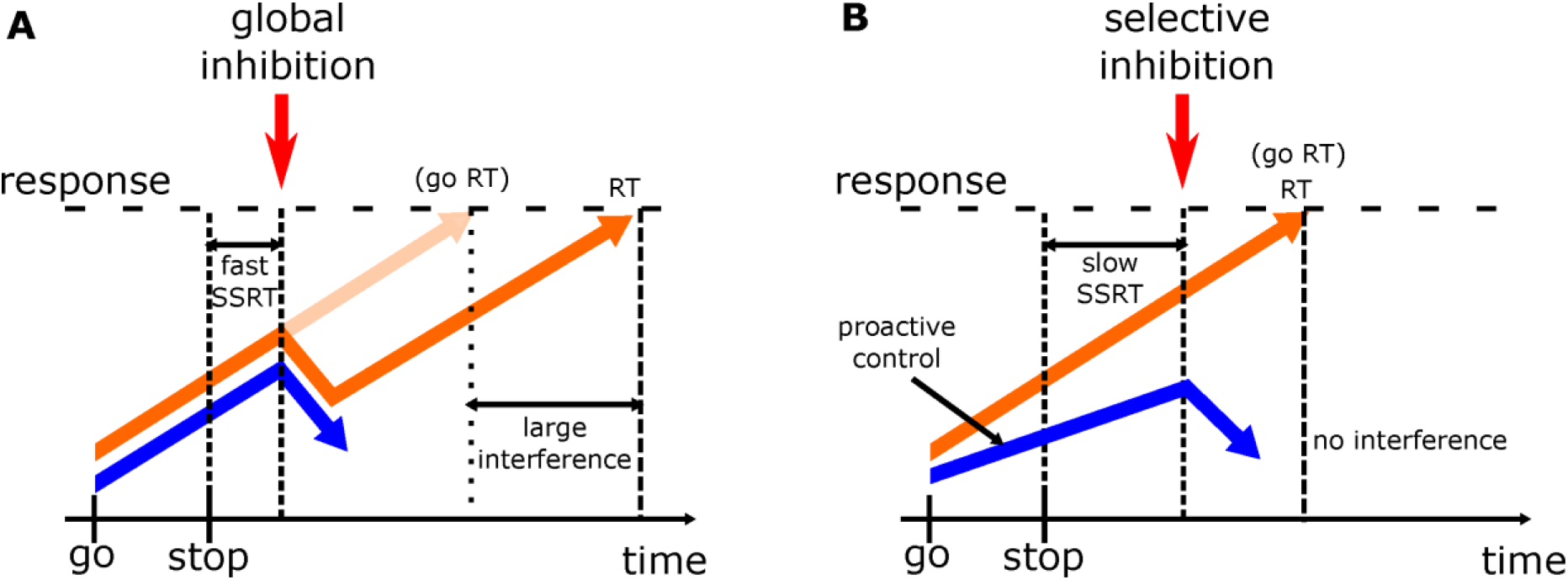
Models for selective stopping in the bimanual-selective stop signal task. The single inhibition model posits that stopping is achieved by a unitary global inhibition mechanism (A), while the dual-mechanism model posits separate global and selective inhibition mechanisms (both A and B). The primary task is to produce go responses with both hands. In case of a stop trial, the response on only one hand needs to be stopped. Orange lines represent activity associated with the responding hand and the blue lines represent activity associated with the stopping hand. **A** represents global inhibition where both responses are inhibited, followed by the restart of the requested response. Global inhibition operates quickly and affects both responses, showing fast stop signal reaction times (SSRT) but delayed reaction times (RT) of continued responses compared to a standard go response (go RT; faint arrow), i.e. an interference effect. **B** represents selective inhibition that is facilitated by proactive control in preparation for the stop signal. In this model stopping operates slower, showing slow SSRTs, but no interference effect, as the responding hand is not affected by inhibition. Note that these models are abstract representations of global and selective inhibition. For simplicity, the effect of proactive control is represented as more gradual activation of the to-be-stopped response (blue arrow), but could also be depicted as a lower starting point for the stop process, hand-specific response threshold modulation, non-stationary activation rate, or any combination of these. The interested reader is directed to more formal models of response inhibition in the uni- or bimanual stop signal tasks (e.g. Boucher et al., 2007; Jahfari et al., 2012; Stuphorn and Emeric, 2012; Macdonald et al., 2014; Schall et al., 2017).

The second model posits distinct global and selective mechanisms (Aron, 2011). Response-specific cues activate selective inhibition via the slow indirect cortico-striatal pathway, prolonging SSRTs, but reducing interference (Aron and Verbruggen, 2008; Claffey et al., 2010). However, others found null or opposite effects for SSRT differences between informative and non-informative cue trials (Smittenaar et al., 2013, 2015; Lavallee et al., 2014; Raud and Huster, 2017; Cirillo et al., 2018). Thus, whether separate mechanisms exist for global and selective inhibition remains unclear (Munakata et al., 2011).

Proactive control can be engaged with cues indicating that a subsequent stop signal, if presented, would require stopping only one component of a multi-effector response. The cues result in anticipatory activation of the stopping network (Chikazoe et al., 2009; Swann et al., 2012) or modulation of attentional and sensorimotor mechanisms (Elchlepp et al., 2016; Langford et al., 2016). In the motor system, down-stream effects are evident in hand-specific modulation of MEPs (Claffey et al., 2010; Jahfari et al., 2010; Cai et al., 2011; Greenhouse et al., 2012), and sensorimotor mu (9-14 Hz) and beta (15-25 Hz) power (Liebrand et al., 2017, 2018), which have been associated with more successful stopping (Mazaheri et al., 2009; Krämer et al., 2011).

We present a multi-modal investigation of stopping, manipulating the demands on proactive control. Inhibition indices were acquired from behavioral, EMG, EEG, and MEP data in uni- and bimanual tasks and we asked whether these dissociate between the global and selective inhibition models. The global account predicts no differences in inhibition indices between tasks, reduced MEPs in an unselected effector during stopping, and interference with ongoing responses during bimanual-selective stopping. The dual-model account predicts slower stopping and reduced interference during bimanual-selective stopping. Further, we expected proactive control to have hand-specific influence on corticomotor excitability and sensorimotor EEG activity before stopping, which in turn would relate to sopping success.

## Materials and methods

### Sample

All participants provided informed consent under a protocol approved by the internal review board of the University of California, Berkeley. Participants were right-handed as confirmed by the Edinburg Handedness Inventory (Oldfield, 1971; missing data for 2 participants who declared right-handedness), had no psychiatric or neurological disorders, and no contraindications to TMS or EEG. The TMS coil was positioned over the EEG cap and electrodes which added an extra 2 cm to the distance between the coil and the skull and necessitated higher TMS intensities to produce MEPs. As such, we restricted testing to individuals with TMS resting motor thresholds (RMTs) below 50% of the maximum stimulator output when not wearing the EEG cap, since the intensity required with the cap in place would have exceeded the maximum stimulator output. Altogether, 33 participants were screened for their stimulation threshold. Ten were excluded due to high thresholds and three withdrew during the TMS thresholding or early in the first session. Data were collected from 20 participants (age 18-29, mean=20.6, sd=2.74, 10 males and 10 females), but because some participants did not complete all sessions or had low data quality in one or more data modalities, the reported analyses include between 17-20 participants (Table 1). Note that the core results did not change if participants with partial data in at least one modality (n=5) were discarded entirely from all analyses.

**Table 1.** Number of participants included in each analysis (out of total n=20) and the reasons for exclusion. The number in the brackets refers to how many participants were excluded due to given reason. *The outlier deviated by 3.4 standard deviations from the group mean.

| Modality | Go trials | Stop trials | Reason for exclusion (# participants excluded) |
| --- | --- | --- | --- |
| Behavior | 19 | 17 | missing unimanual session (1)<br>stop accuracies below 30% (2) |
| EMG | 19 | 19 | missing unimanual session (1) |
| MEP | Unimanual: 18<br>Bimanual: 20 | 17 | missing unimanual session (1)<br>less than 10 post-stop MEPs (2)<br>outlier* in unimanual pre-go MEPs (1) |
| EEG | Unimanual: 17<br>Bimanual: 20 | 17 | missing unimanual session (1)<br>technical EEG problems in unimanual session (1)<br>artifacts due to TMS coil in unimanual session (1) |

### Task and procedure

The participants performed two versions of the stop signal task – unimanual and bimanual – with response-specific cues (Figure 2). The tasks were performed in separate experimental sessions and the order of the tasks was counterbalanced across participants. Participants sat at a viewing distance of about 50 cm in front of the computer screen (refresh rate 80 Hz), hands laid palms down on pillows placed on a table in front of them so that the thumbs (responding effectors) rested on a keyboard used to collect responses. The experimental protocol was implemented with Psychtoolbox 3.0.

**Figure 2.**
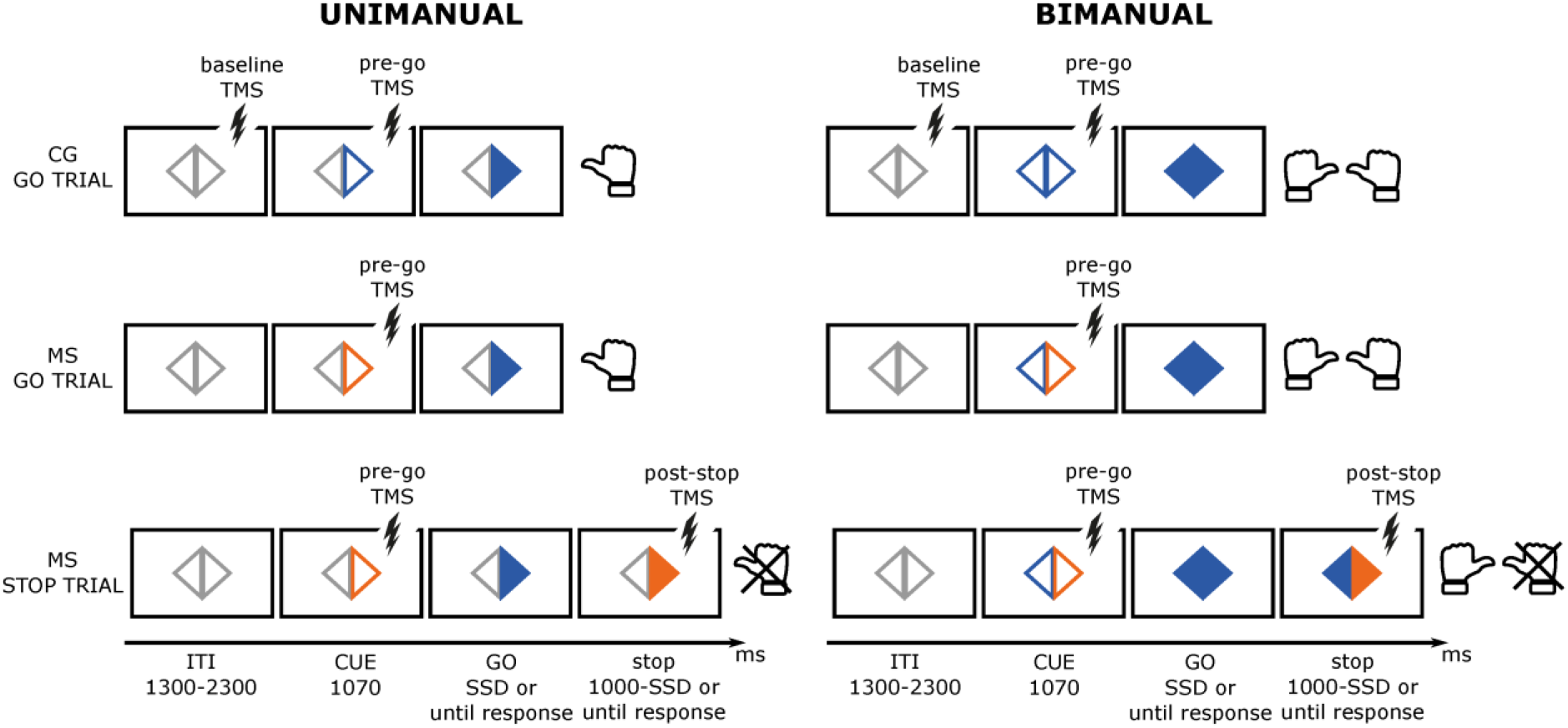
Unimanual and bimanual-selective stop signal tasks. The baseline TMS pulses were given 200 ms prior to the cue, pre-go TMS pulses 100 ms prior to the go signal and post-stop TMS pulses 150 ms after the stop signal. TMS was applied on approximately 27% of trials. All trial types were divided into go/stop and left/right trials (only go/stop right trials shown on figure). Responses were given as thumb presses. CG – certain go; MS – maybe stop.

Two hollow triangles were present on the screen at all times and served as the fixation point during the inter-trial intervals. In the unimanual task, the change of the outline color (from gray) of a single triangle signaled the participant to prepare a response with either the left or right thumb, depending on which triangle changed color. The specific color of the outline of the triangle cued the participant whether the given trial was a certain go trial (CG) or whether it may be a stop trial (MS). The triangle then filled with color (the go signal) to signal participants to execute their prepared response. On a fraction of trials, the triangle changed color again (the stop signal) signaling participants to stop their prepared or initiated response.

The bimanual task was similar, except that both triangle outlines changed color after the inter-trial interval, cueing participants to prepare thumb responses with both hands. In the CG condition, both triangle outlines were of the same color, whereas in the MS condition the triangle outlines were of different colors with one color indicating which hand’s response may have to be stopped. On go trials, both triangles filled with the same color, instructing the participants to produce a simultaneous response with the left and right hand. On stop trials, only the previously cued triangle changed color again, instructing the participants to stop the cued hand while continuing the response with the other hand. The colors for go and stop stimuli (and corresponding CG and MS cues) were blue and orange shown on a gray background, with the color mapping counterbalanced between participants.

Each trial started with the presentation of the cue for 1070 ms, indicating the response hand(s) and possibility of stopping. The cue was followed by the go signal. On go trials, the go signal remained visible until 100 ms after a response was registered or until the end of a response window of 1000 ms. On stop trials, the relevant triangle changed to the stop signal color after the stop signal delay (SSD). The SSD was adjusted according to a staircase procedure separately for stop left and stop right hand trials. The SSD increased in steps of 47 ms if stopping was successful on the previous trial, and otherwise decreased by 47 ms if stopping failed. The stop signal was presented until 100 ms after a response (unsuccessful stop) or until the end of the response window (successful stop). The inter-trial interval varied between 1300-2300 ms.

In the unimanual task, the trials were divided equally into left and right hand trials. In the bimanual task, all go trials were the same (bimanual go), but stop trials were equally divided into stop left and stop right hand trials. All trials were further divided into those with and without TMS. Each task consisted of 200 CG-go trials, 680 MS-go trials, and 340 stop trials (see Table 2 for the number of trials included in each analysis modality). The stop signal was presented on 33% of the MS trials and 26% of all trials. To prevent premature responding during the cue period, 88 catch trials (20 CG and 68 MS, corresponding to 10% of go trials) were added, in which the go signal was not presented after the cue. The total number of trials amounted to 1308 per task, divided into 10 blocks (each lasting about 8 minutes) with pauses of self-determined durations between the blocks. The order of trials in each block was pseudorandomized.

**Table 2.** Number of trials (standard deviations in brackets) per condition. Total refers to the total number of trials presented across all modalities and the rest refer to the mean number of trials per condition, participant, and analysis modality (standard deviations in the brackets) that were included in the analysis after data cleaning. Baseline MEP trials and catch trials are omitted from the table. The MEPs are further divided according to whether the right hand was selected or unselected for the response (unimanual task) or selected for stopping or responding (bimanual task). *Maybe stop pre-go TMS pulses were given both in go and stop trials with a ratio of 30/10 to prevent the prediction of the upcoming stop signal from application of the pre-go pulse. N.a. – not analyzed.

| GO TRIALS |  |  |  |  |  |  |  |
| --- | --- | --- | --- | --- | --- | --- | --- |
| UNIMANUAL |  |  |  |  | BIMANUAL |  |  |
|  | certain go |  | maybe stop |  | certain go | maybe stop |  |
| Total | 200 |  | 680 |  | 200 | 680 |  |
| Without TMS |  |  |  |  |  |  |  |
| BEH | 93 (14) |  | 596 (58) |  | 91 (8) | 587 (32) |  |
| EMG | 83 (16) |  | 564 (80) |  | 85 (10) | 557 (47) |  |
| EEG | 73 (16) |  | 466 (70) |  | 69 (12) | 447 (53) |  |
| With TMS (pre-go)* |  |  |  |  |  |  |  |
|  | selected | unselected | selected | unselected | selected-go | maybe stop | selected-go |
| MEP | 29 (6) | 32 (6) | 32 (4) | 32 (5) | 61 (11) | 32 (5) | 31 (5) |

**Table 2.** Number of trials (standard deviations in brackets) per condition.
| STOP TRIALS |  |  |  |  |  |  |  |  |
| --- | --- | --- | --- | --- | --- | --- | --- | --- |
| UNIMANUAL |  |  |  |  | BIMANUAL |  |  |  |
|  | successful |  | unsuccessful |  | successful | unsuccessful |  |  |
| Total | 340 |  |  |  |  | 340 |  |  |
| Without TMS |  |  |  |  |  |  |  |  |
| BEH | 89 (12) |  | 86 (14) |  | 89 (9) | 86 (8) |  |  |
| EMG | 80 (18) |  | n.a. |  | 84 (10) | n.a. |  |  |
| EEG | 66 (15) |  | 65 (15) |  | 73 (12) | 66 (9) |  |  |
| With TMS (post-stop) |  |  |  |  |  |  |  |  |
|  | stop | unselected | stop | unselected | stop | response | stop | response |
| MEP | 20 (7) | 33 (8) | n.a. | n.a. | 24 (7) | 20 (7) | n.a. | n.a. |

Participants received instantaneous feedback if they did not respond within the response window of 1000 ms in go trials or if their response was longer than 800 ms to prevent strategic slowing due to stop signal expectancy. A red exclamation mark appeared on the screen in the bimanual task if the difference between left and right hand responses exceeded 70 ms. Finally, block-wise feedback was given, prompting the participants to be faster if the reaction times of the previous block on average were slower than 600 ms, or to be more accurate if stopping accuracy fell under 45%.

### TMS procedure

Single TMS pulses were delivered with a 70 mm figure-of-eight coil driven by a Magstim 200-2 magnetic stimulator. The coil was positioned tangentially over the left motor cortex in the posterior-anterior direction and the position was optimized to elicit MEPs in the right abductor pollicis brevis (APB) muscle. The stimulation intensity was set to 115% of the RMT, defined as the intensity where 5 out of 10 consecutive pulses elicited a MEP with an amplitude of at least 50 μV. This threshold was determined after EEG preparation to account for the increased distance between the coil and the scalp due to the electrodes. The coil rested on a plastic spacer placed between the coil and the EEG-cap to reduce mechanical EEG artifacts due to the pressure of the coil (Ruddy et al., 2018).

TMS pulses during the task were administered pseudorandomly between all trials with an inter-stimulus-interval between 4 and 54 seconds. The total number of pulses per session was 360. However, some pulses were omitted (~12% per session) when a pulse was scheduled by pseudorandomization with an inter-stimulation interval <4 sec or when the participant terminated the trial before the scheduled pulse time. To establish a baseline measure of corticospinal excitability, on 40 CG-go trials TMS pulses were scheduled 200 ms prior to the cue (Figure 2). The pre-go pulses were scheduled 100 ms before go onset (corresponding to 970 ms after the cue) on 80 CG and 80 MS trials (further divided into 60 go and 20 stop trials). The post-stop pulses were scheduled at 150 ms following the stop signal on 160 trials, as we expected inhibition to be present around this time in stop trials, based on previous TMS and EMG results (van den Wildenberg et al., 2010; Macdonald et al., 2014; Raud and Huster, 2017; Atsma et al., 2018).

### EEG and EMG data acquisition

EMG was recorded from the left and right hand abductor pollicis brevis with bipolar Ag/Cl surface electrodes (Delsys, Inc.) placed in parallel to the belly of the muscle. The ground electrode was placed over the ulna of the right elbow. The data were recorded with a sampling rate of 4000 Hz and bandpass filtered between 20-450 Hz during data acquisition. EEG was recorded with the Active2 system from 64 channels (BioSemi) at a sampling rate of 2048 Hz (low-pass filter 417 Hz). Additional electrodes were placed on the nose-tip, mastoids, next to the outer canthis and below the right eye.

### Data preprocessing and feature extraction

#### Behavior

The following variables were determined for trials without TMS pulses: go accuracy, go reaction time (RT), stop accuracy, stop signal delay (SSD), stop signal reaction time (SSRT), unsuccessful stop trial RT. Additional variables available only for stop trials of the bimanual task included the responding hand RT and behavioral interference, calculated as the difference between the responding hand RT in successful stop trials and the MS go RT. In correct go trials, very fast (<100 ms) go RTs were discarded from the RT calculation (0.3 %). In the bimanual task, the RT was obtained by averaging over both hands. The SSRT was calculated by the integration method with replacements for go omissions and including go errors (Verbruggen et al., 2019). The data from left and right hand trials were averaged for statistical analysis.

#### MEPs

MEP amplitudes were measured as the peak-to-peak amplitude in the EMG response in the 10-100 ms window following the TMS pulse. Trials were rejected when the average rectified EMG activity 200 ms prior to the pulse was over 10 μV. Additionally, individual trial data were visualized, and trials with EMG activity just before or at the time of the TMS pulse were discarded (26% on average across all conditions). MEP amplitudes covary with background muscle activity, which in turn may be affected by task conditions (Figure 4A). For this reason, we residualized the MEP amplitudes for the pre-pulse RMS activity (Figure 4B). We extracted the average RMS 50 ms before each TMS pulse and fit a regression model for each participant, predicting the MEP amplitudes from the pre-pulse RMS. The residuals of these models were distributed into conditions and were normalized by dividing by the residualized baseline MEP amplitudes (shifted by a positive coefficient to prevent distortions of the data from division with negative residuals). Thus, the statistical analysis was performed on the MEP amplitudes representing the ratio to the baseline after controlling for the differences in the pre-pulse background muscle activity.

#### EMG

EMG was downsampled to 512 Hz, incorporating a polyphase anti-aliasing filter. Continuous data were epoched relative to the cue onset with an epoch length of −300 to 2600 ms. A moving average procedure with a window size of +/−5 datapoints was applied to the root mean square of the signal, and this was normalized by dividing each datapoint with the average baseline period of −300 to 0 ms relative to the cue onset. The epochs of all trials were concatenated for each participant and hand, and the data was z-scored across all conditions. The data was then re-epoched into the conditions of interest, with an epoch length of −300 to 1800 ms for cue-locked activity, and −300 to 600 ms for stop-locked activity.

An automatic artifact rejection and EMG-burst identification algorithm was applied to the epochs, where an EMG burst in a given trial was determined whenever the z-scored RMS data exceeded a threshold of 1.2. This threshold was chosen by visual inspection of several datasets and was held constant for all datasets. Trials were rejected automatically if the average rectified baseline activity exceeded 10 μV or if an EMG burst was detected prior to the go stimulus onset. All trials were further visualized and trials with excessive tonic muscle activity were discarded. Altogether, 3.75% trials per condition were discarded on average at this step.

The following dependent variables were extracted: 1) prEMG burst frequency: the percentage of successful stop trials with EMG activity above threshold relative to the total number of successful stop trials; 2) EMG onset latency: the latency of the first data-point exceeding the EMG threshold; 3) EMG peak latency: the time point at which the EMG activity reached its maximum. Onset latencies were calculated relative to the go-signal onset, while the peak latency of the prEMG was calculated relative to the stop-signal onset. Trials in which the amplitude or latency measure exceeded +/−2.5 standard deviations of each participant’s mean were discarded from the analysis, as well as trials where prEMG peaked before the stop signal or more than 500 ms after the stop signal, as these were likely not related to stop signal processing (altogether 2.54% per condition on average). To capture potential interference in the responding hand EMG in the bimanual task at the single trial level, the Δpeak was calculated by subtracting the prEMG peak latency of the stopped hand from the EMG peak latency of the responding hand (see Figure 3A for schematics).

**Figure 3.**
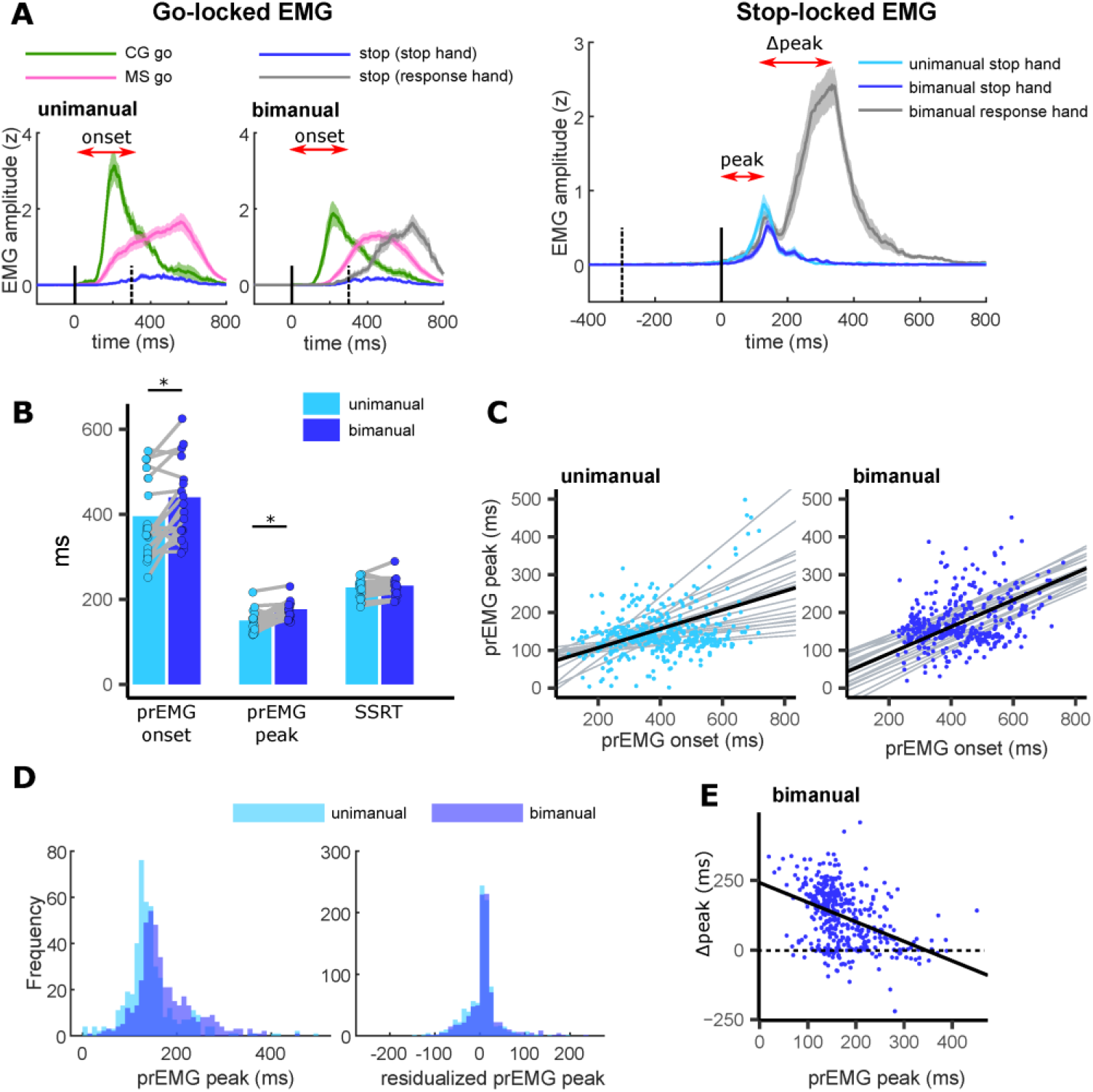
**A**. EMG time courses time-locked to the go signal (left panels) and the stop signal (right panel). The red arrows reflect the parametrization of prEMG onset and peak, and Δpeak for the bimanual task. The solid vertical lines represent the time-locking event and the dashed vertical lines represent the average SSD. **B**. Average prEMG onsets, peaks, and SSRTs across participants. The dots represent individual participants and the gray lines connect the same participants in the uni- and bimanual task. Asterisks indicate task differences with BF > 3. **C**. Single trial correlations between prEMG onset and prEMG peak. The black lines indicate the fixed effects of prEMG onset on prEMG peak, and gray lines indicate the random effects (intercept and slope) for each participant. **D**. Histograms of prEMG peak latencies across all trials and participants before (left panel) and after (right panel) residualizing for the prEMG onset. **E.** Single trial correlation between prEMG peak and Δpeak latency in the bimanual task. The horizontal dashed line marks zero Δpeak; data points in the vicinity of the line correspond to trials with small or no interference. CG – certain go; MS – maybe stop.

**Figure 4.**
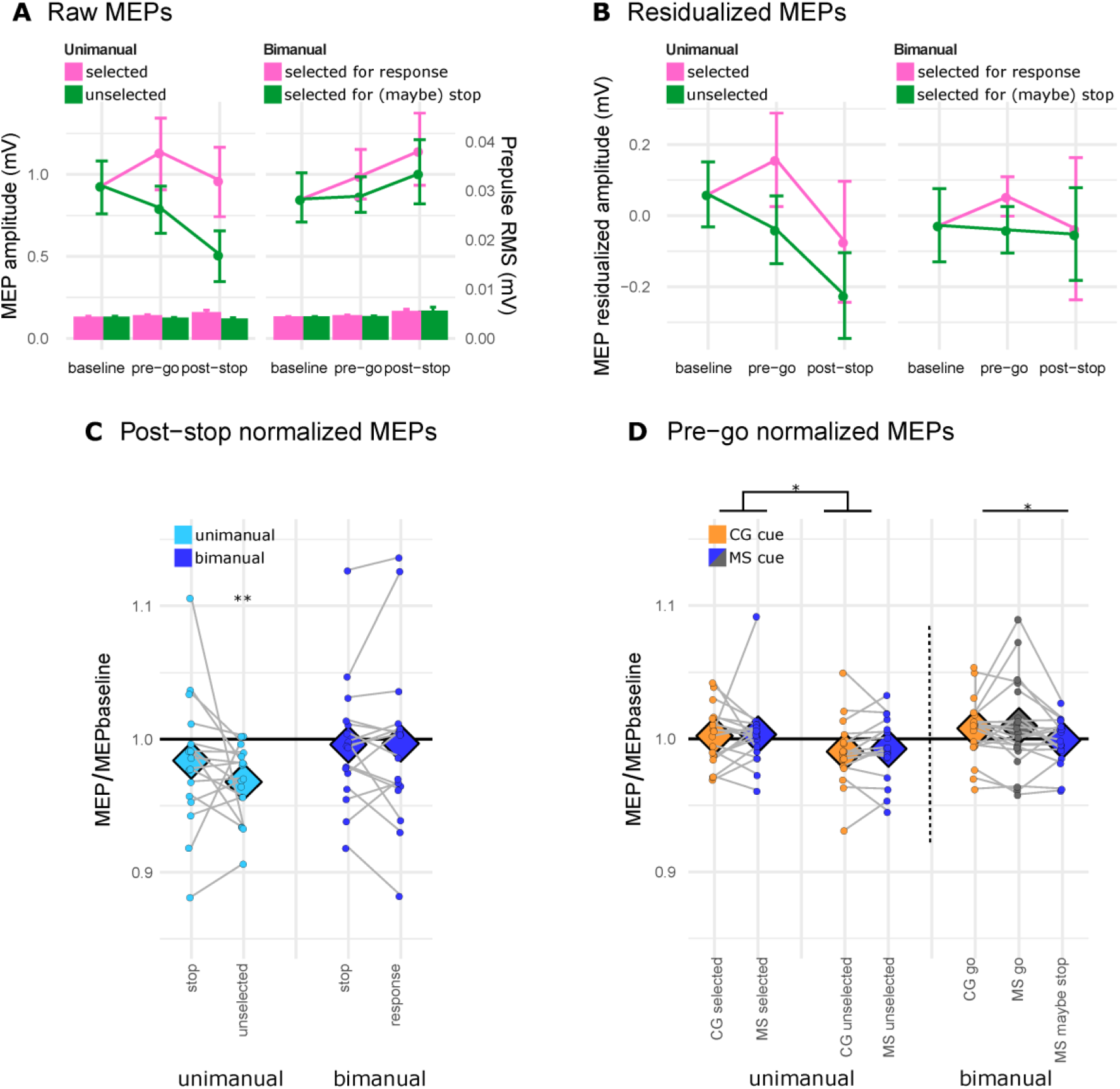
**A-B** Raw MEPs (A) and residualized MEP amplitudes after fitting a regression on a single participant level, predicting the raw MEP amplitude from the EMG root mean square activity 50 ms prior to the TMS pulse (B). The pre-go MEPs are averaged over certain go and maybe stop cues for A and B. The bars in A represent the pre-pulse RMS activity (associated with the right y-axis). **C-D** Condition specific (residualized) MEPs as a ratio to the baseline MEPs, measured after the stop signal (C) and before the go signal (D). The diamonds represent average values across participants, the dots represent single participants, and the gray lines connect the same participants in selected conditions. The dashed vertical line on (C) represents the separation between unimanual and bimanual

#### EEG

EEG was analyzed with *eeglab* v14.1.2 (Delorme and Makeig, 2004). Trials with TMS pulses were discarded from the analysis, and the data from −5 to 500 ms relative to each TMS pulse were replaced using a cubic interpolation to avoid possible filtering artefacts that could contaminate the trials directly before or after the pulse. The data were low-pass filtered using a hamming window filter with a 6db cut-off at 50 Hz (transition bandwidth 20 Hz), resampled to 512 Hz, and high-pass filtered with a 6db cut-off at 0.1 Hz (transition bandwidth 0.8 Hz). The data were epoched relative to the cue onset, and trials with artefacts were rejected manually by visual inspection (21% on average). At this step, noisy channels were identified and temporarily discarded (1.16 electrodes or 1.82% on average). An independent component analysis was run on the remaining data and artifactual components reflecting eye-movements and muscle activity were identified and rejected semi-automatically (5.49 components or 8.57% per dataset on average) using the *eeglab* plugin SASICA (Chaumon et al., 2015). An interpolation procedure was used to replace data from previously identified noisy electrodes. Lastly, to reduce the volume conduction in the scalp EEG, a current source density interpolation was estimated by a spatial Laplacian transformation using the current density toolbox for Matlab (Kayser and Tenke, 2006).

For time-frequency analyses, the data were segmented into condition-specific epochs of −1200 to 3000 ms. Time-frequency decompositions for 1-40 Hz were calculated using Morlet wavelets with the number of cycles linearly increasing from 1 to 20. The cue-locked activity was normalized by dividing the power by the average of the frequency-specific baseline between −400 to −100 ms relative to cue-onset. The stop-locked activity was similarly normalized to the average of −400 to −100 ms before the stop signal.

Frontal beta (13-20 Hz) activity was extracted from 100 to 150 ms after the stop signal, averaged over the frequencies and electrodes of interest: left prefrontal (F5, F3, FC5, FC3), frontocentral (F1, Fz, F2, FC1, FCz, FC2) and right prefrontal (F4, F6, FC4, FC6). Electrodes and frequencies were determined based on previous studies showing frontal beta effects in the stop signal task. Beta frequencies vary considerably between different studies, and we chose this range of frequencies based on a recent study that reported a beta power increase in frontal scalp EEG electrodes, presumably reflecting the activity of the stopping network in the prefrontal cortex (Wagner et al., 2018). The time-window was selected on the rationale that brain activity prior to the peripheral inhibition latency (i.e. prEMG latency at ~150 ms) would be the best indicator of inhibitory control.

Sensorimotor mu (9-14) and beta power (15-25 Hz) were extracted from the cue-locked activity in pre-go (−200 to 0 ms) and post-go (100 to 200 ms) time-windows on go trials, and post-go time-window (100 to 200 ms) on stop trials. The pre-go time-window was selected so that it would match the timing of the pre-go TMS pulses (−100 ms relative to go) to allow for comparisons between the EEG and MEP results. The post-go time window was selected so that it would correspond to the time after the initial sensory processing and before response execution (i.e. EMG onset). Frequencies and electrodes of interest were determined based on previous studies (Krämer et al., 2011; Liebrand et al., 2018), corresponding to left (C5, C3, CP5, CP3) and right (C4, C6, CP4 and CP6) sensorimotor regions. The electrodes and frequencies used for this analysis are different from those used for the stop-locked activity analysis because earlier work suggests that motor activity in sensorimotor cortices differs functionally from the frontal beta oscillations typically associated with response inhibition.

### Experimental design and statistical analysis

Group-level statistical analyses were performed in the Bayesian framework using repeated measures ANOVAs (rmANOVA) and paired samples t-tests with uniform priors. The Bayes factors (BF) were calculated with the package BayesFactor in R. The BF’s reported here represent the likelihood for the model with a main effect of interest compared to the model with random participant intercepts. For the interaction terms, the BF’s represent the likelihood of the interaction model against all models that include the specific terms that form the interaction (calculated with R package bayestestR). The BF values below and above one indicate the likelihood for the null or alternative hypothesis, respectively, and BF=1 means that the null and alternative hypothesis are equally likely.

Normality of all the variables was tested by Shapiro-Wilk tests. In cases of deviations of normality, the analysis was repeated on the log-transformed variables. Given that this analysis did not lead to any deviations from the main conclusions, we visualize and report the statistics of non-transformed data to simplify the interpretation.

The statistical design is described in more detail in reference to the three sets of predictions outlined in the introduction.

#### 1) Do indices of inhibition in the uni- and bimanual stop signal task dissociate between the two models of inhibition?

The SSRTs and prEMG latencies of the two tasks were compared by means of paired samples t-tests. The post-stop MEPs were tested with a one-way rmANOVA (levels unimanual stop hand, unimanual unselected hand, bimanual stop hand, bimanual response hand) and followed up by pairwise t-tests. The frontal beta power was tested using a three-way rmANOVA with factors SUCCESS (successful, unsuccessful), TASK (unimanual, bimanual) and electrode LOCATION (frontal contralateral, central, frontal ipsilateral). ‘Contralateral’ refers to the electrodes positioned contralateral to the stopped hand (and thus the electrodes placed ipsilateral to the hand still responding in stop trials of the bimanual task).

Given that the prEMG analysis indicated differences both between go-locked prEMG onset latencies and stop-locked prEMG peak latencies, we conducted an additional analysis to dissociate whether these effects were driven by the go process, stop process, or the interaction of the two. First, the prEMG peak latency analysis was repeated after residualizing for the prEMG onsets. Secondly, we tested the relationship between the peaks and onsets on a single trial basis using linear mixed effects regression models to control for the influence of individual participants. The regressions were fitted with the R package *nlme*. Given the individual differences in the general response speed, we considered the model with random intercepts as the baseline model and tested the improvement in model fit after including random slopes with chi-square tests. Both unstandardized and standardized coefficients are reported, marked as b and b_std_, respectively.

Lastly, we examined the EMG measure of interference (Δpeak) in detail and tested whether it is associated with stopping latency (prEMG latency) at the single trial level. This was done using the linear mixed effects models as described above.

#### 2) Does motor preparation influence subsequent inhibition success?

Given the putative roles of sensorimotor mu and beta activity in action preparation (Neuper et al., 2006; Brinkman et al., 2016), we hypothesized that mu and beta activity before the stop signal would predict stopping success. Condition differences were tested using rmANOVAs incorporating the SUCCESS (successful, unsuccessful) of stopping and electrode LOCATION (contralateral and ipsilateral hemisphere relative to the stopped hand) as factors. This was done separately for the tasks, as the differences in the go instructions did not allow for a balanced statistical comparison regarding the hemispheric effects.

#### 3) Is motor preparation affected by proactive control?

In the current study, proactive control is relevant in conditions in which the participants are informed that a forthcoming trial may require stopping. We tested the effect of proactive control on response preparation by comparing trials with maybe stop (MS) and certain go (CG) cues. The effects of the cue on the go RTs were tested with rmANOVAs including the factors CUE and TASK. MEPs were tested separately in each task. In the unimanual task, we performed a two-way rmANOVA with factors CUE and HAND (selected, unselected). In the bimanual task, we performed a one-way rmANOVA with a single combined CUE-HAND factor with levels CG, MS go (i.e. respond right, maybe stop left), and MS maybe stop (i.e. maybe stop right, respond left). The effects of the cue on sensorimotor mu and beta power were also tested separately for each task using rmANOVAs. Here, we additionally tested the difference between pre-go and post-go activity, to determine whether the effects were evident in the cue-delay period before the go stimulus presentation. In the unimanual task, the factors were CUE, electrode LOCATION, and TIME. In the bimanual task, the levels for a combined CUE-LOCATION factor were CG bilateral, MS contra-go, MS contra-stop. In the CG-bilateral, activity was averaged over left and right hemisphere electrodes given the bilateral response, while the MS contra-go refers to the electrodes contralateral to the responding hand, and MS contra-stop refers to the electrodes contralateral to the hand that may need to be stopped.

## Results

### 1) Do inhibition indices in the uni- and bimanual stop signal task dissociate between the two models of inhibition?

#### No differences between the SSRTs of the uni- and bimanual tasks

Task performance was in accordance with expectations based on the horse race model, with stopping accuracies at 51% in both tasks (see Table 3 for all behavioral results) and faster unsuccessful stop RTs than go RTs in all individuals. The SSRTs were similar between the two tasks (BF = 0.33; Figure 3B).

**Table 3.** Means and standard deviations of the behavioral parameters in both tasks for trials without TMS. CG – certain go; MS – maybe stop; RT – reaction time; SSD – stop signal delay; SSRT – stop signal reaction time. *In the unimanual task, errors refer to the trials where the response was executed using the wrong hand; in the bimanual task, errors refer to trials where a response was executed with only one hand.

|  | Unimanual | Bimanual |
| --- | --- | --- |
| **Go trials** |  |  |
| CG accuracy (%) | 94.71 (6.21) | 95.62 (3.70) |
| omissions | 1.40 (2.39) | 0.85 (1.63) |
| errors* | 0.20 (0.66) | 0.39 (0.74) |
| premature | 3.70 (4.57) | 2.18 (2.36) |
| asynchronous (>70 ms) | - | 0.97 (1.22) |
| MS accuracy (%) | 97.18 (3.05) | 94.73 (3.28) |
| omissions | 2.01 (2.65) | 0.91 (1.79) |
| errors* | 0.31 (0.42) | 1.33 (1.44) |
| premature | 0.51 (0.63) | 0.31 (0.30) |
| asynchronous (>70 ms) | - | 2.72 (1.71) |
| CG RT (ms) | 364 (79) | 388 (86) |
| MS RT (ms) | 518 (98) | 532 (78) |
| CG asynchrony (ms) | - | 14 (3) |
| MS asynchrony (ms) | - | 15 (4) |
| **Stop trials** |  |  |
| Stop accuracy (%) | 50.85 (4.08) | 50.94 (3.04) |
| Unsuccessful stop RT (ms) | 456 (93) | 486 (72) |
| SSD (ms) | 307 (92) | 307 (88) |
| SSRT (ms) | 228 (23) | 232 (22) |
| Response hand RT (ms) | - | 656 (72) |
| Behavioral interference (ms) | - | 113 (36) |

#### Delayed prEMG peaks in the bimanual task, possibly resulting from dependencies between go and stop processes

Successful stopping was defined as trials in which a key press was not detected. However, on approximately 30% of these trials (an average of 24 trials (sd = 14) in the unimanual and 22 (sd = 10) in the bimanual task for each participant), EMG activity was detected in the stopped hand, or what we refer to as prEMG (Table 4). The onset of prEMG relative to the go signal provides an index of the speed of the go process on these trials, while the latency of the decline of this signal (i.e. the prEMG peak latency) relative to the stop signal provides an index of the speed of the inhibitory process (Figure 3A). The bivariate correlations between the SSRT and prEMG peak latency were 0.52 in the unimanual (BF = 2.99) and 0.14 in the bimanual task (BF = 0.57).

**Table 4.**
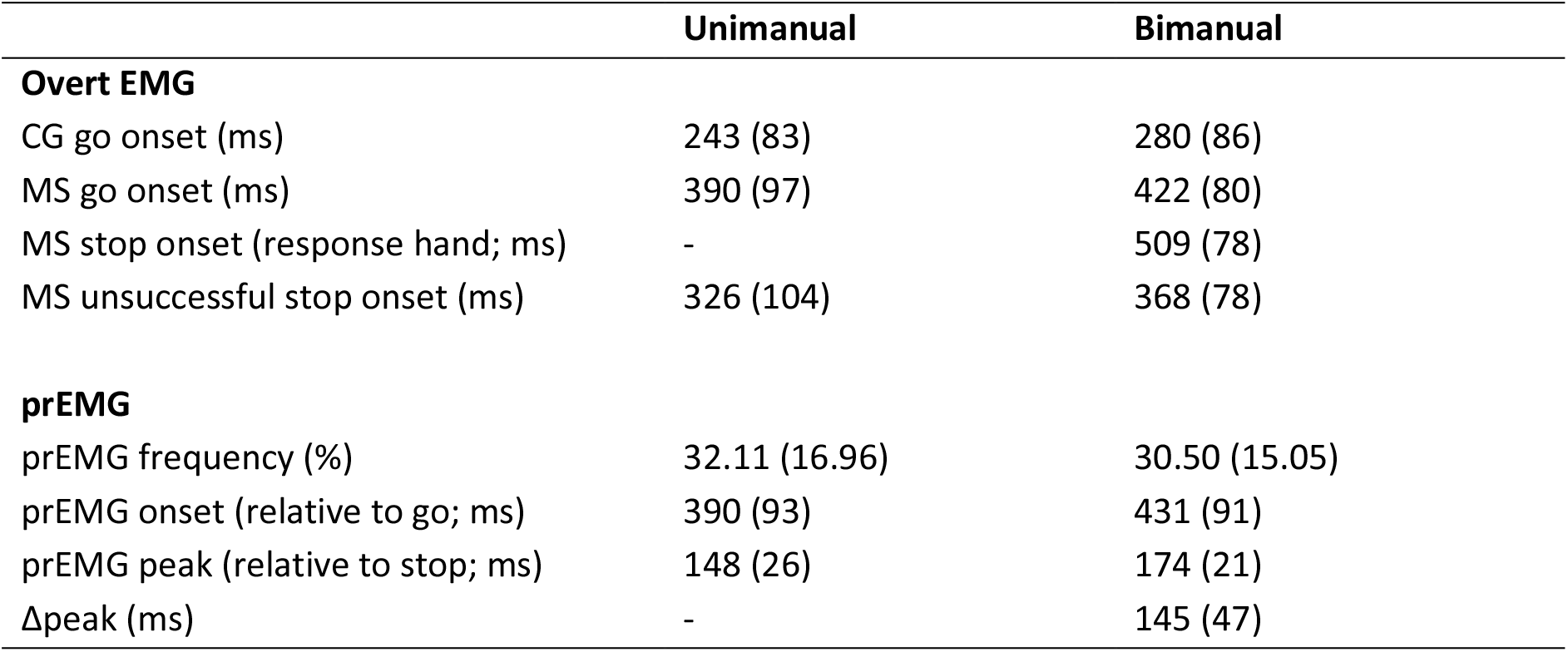
Mean and standard deviation of overt EMG (where button presses were registered) and prEMG in successful stop trials (where no button press was registered). CG – certain go; MS – maybe stop.

The prEMG peak latency was later in the bimanual task compared to the unimanual task (BF = 233.69, Figure 3A, 3B). However, prEMG onset was also significantly later in the bimanual than in the unimanual task (BF = 6.61). When comparing the EMG onsets in go and stop trials within a single rmANOVA with factors TRIAL (go, stop) and TASK (unimanual, bimanual), we again find an effect of the TASK (BF = 133.45) but no effects of the TRIAL (BF = 0.26) nor interaction of the two (BF = 0.31), suggesting that the task differences are not specific for stopping. The delayed go and stop processes in the bimanual task further raise the possibility that there are dependencies between the two processes, and that these dependencies may differ between the tasks. This is important for the evaluation of the latency differences between the tasks, as dependencies would violate assumptions of the horse race model that are necessary for a reliable estimation of SSRTs. We explored this possibility at the single trial level on a post-hoc basis using mixed linear regression models, where prEMG peak latency was predicted by the prEMG onset latency and task type, with participants’ intercepts and slopes as random effects. We fit models predicting prEMG peaks from onsets separately in both tasks (Figure 3C). In the unimanual task, the fixed effect indicated a positive relationship between the peaks and onsets (b = 0.25, 95% CI = 0.15, 0.35, t(430) = 4.82, p < 0.001, b_std_ = 0.54) and considerable variability in the slopes across participants (sd = 0.19, 95% CI = 0.13, 0.29). There was a significant improvement in model fit when including the variability of slopes, compared to the random intercept model (*χ*^2^(1) = 62.75, p < 0.001), indicating that the relationship between peaks and onsets was relatively heterogeneous among the participants (Figure 3C, left panel). In the bimanual task, the fixed effects also indicated a positive relationship between peaks and onsets (b = 0.36, 95% CI = 0.29, 0.43, t(404) = 10.01, p < 0.001, b_std_ = 0.67), but smaller variability in the slopes (sd = 0.08, 95% CI = 0.02, 0.28). The improvement in model fit was not significant after including variable slopes (*χ*^2^(1) = 1.53, p = 0.465), suggesting that the association between the onsets and slopes were relatively similar across participants (Figure 3C, right panel).

Given the positive association between prEMG onsets and peaks, we next asked if the task differences in prEMG peak latencies could be explained by the differences in the onsets. We tested the differences between the peaks again, but this time after residualizing for the differences in onset times. The histograms for prEMG peaks before and after residualization (Figure 3D) reveal that the initial differences between the tasks were greatly diminished after residualization. In fact, this analysis showed that the data is just as likely in the case of the null hypothesis as the alternative hypothesis (BF = 1.00).

#### Prevalent and variable interference in the bimanual task

The RTs of the responding hand on bimanual successful stop trials were delayed compared to go RTs (MS cue), giving rise to a mean behavioral interference effect of 113 ms. This interference was also evident in the EMG of the continuing response hand (grey lines on Figure 3A, right panel). There was a halt in the progression of the EMG activity at around 170 ms, followed by the recovery of the EMG activity leading to the key press.

EMG allows for the investigation of the interference effect and its associations at a single trial level. To achieve this, we extracted the Δpeak (the difference between responding hand EMG peak and stopped hand prEMG peak) from all trials where prEMG was detected. The average Δpeak on the group level was 145 ms, but varied within trials between −219 and 459 ms, with a negative Δpeak indicating that the EMG in the responding hand peaked before the prEMG in the stopped hand. At the group level, the Δpeak correlated positively with the behavioral interference (r = 0.85, BF = 1278.90). Further, the single trial Δpeak correlated negatively with prEMG peak latency, as tested by the regression model with random intercepts (Figure 3E; b = −0.70, 95% CI = −0.84, −0.56, t(404) = −9.69, p < 0.001, b_std_ = −0.42), indicating that later prEMG peaks occurred on trials with reduced interference. This relationship remained similar after residualizing the peaks by prEMG onset. The large single-trial variability in the interference effect together with its associations with the stopping latency suggests that successful stopping in the bimanual task may include a mix of trials with different behavioral strategies, which also play a role in regulating inhibition speed.

#### Post-stop MEPs are reduced in the unimanual unselected hand, but do not differ between the uni- and bimanual tasks for the stopped hand

Resting motor thresholds were 70% (sd = 5.67) and 71% (sd = 6.86) of maximum stimulator output in the uni- and bimanual task respectively, and the raw baseline MEP amplitudes were similar between the two tasks (unimanual: 0.93 (0.71) mV; bimanual: 0.88 (0.69) mV; BF = 0.26). The post-stop MEPs, probed by a TMS pulse applied over left motor cortex 150 ms after stop signal presentation, were measured on trials where either the left or right hand was stopped, resulting in four different post-stop MEP conditions: unimanual stop (i.e. MEPs measured from the right hand on right hand stop trials), unimanual unselected (i.e. MEPs measured from the right hand on left hand stop trials), bimanual stop (i.e. stop right, respond left), and bimanual response (i.e. stop left, respond right; Figure 4C). As predicted by the global inhibition model, MEPs in the unimanual unselected condition were suppressed compared to baseline (BF = 134.56), and all other conditions did not differ from baseline (BFs <= 0.56). The omnibus one-way ANOVA indicated considerable differences between the conditions (BF = 565.14). The pairwise post-hoc tests indicated that the unselected hand MEPs in the unimanual task differed from the stopped hand in the bimanual task (BF = 1.69). Given that the dual-mechanisms account predicts slower inhibition in the bimanual task, we expected that the MEP suppression would be weaker in the bimanual task at the specific time-point when the pulse was applied (i.e. 150 ms after the stop signal). Critically, there was no evidence for MEP amplitude difference between the unimanual stop and bimanual stop conditions (BF = 0.46), nor was there evidence for differences between the stopped and responding hand MEP amplitudes in the bimanual task (BF = 0.25).

#### No differences in frontal beta between uni- and bimanual tasks

Frontal beta may be a cortical index of response inhibition, and may therefore differentiate between the tasks. Specifically, we looked at the early differences prior to the prEMG peak latency, expecting stronger beta in successful than unsuccessful trials. In addition, lower beta power in the bimanual task may be indicative of a slower and more selective inhibitory mechanism. An increase in beta power was observed in all stop trials, though it appeared to be strongest at later time-points (Figure 5) and was close to zero in the time-window of 100-150 ms after the stop signal (avg. beta over all conditions = 0.08 dB (sd = 0.42), BF = 0.32). In contrast to our predictions, there were no significant differences between the levels of TASK (BF = 0.17) nor LOCATION (BF = 0.06). There was weak evidence for the effect of SUCCESS (BF = 1.42), while the interaction between SUCCESS and TASK was inconclusive (BF = 1.03).

**Figure 5.**
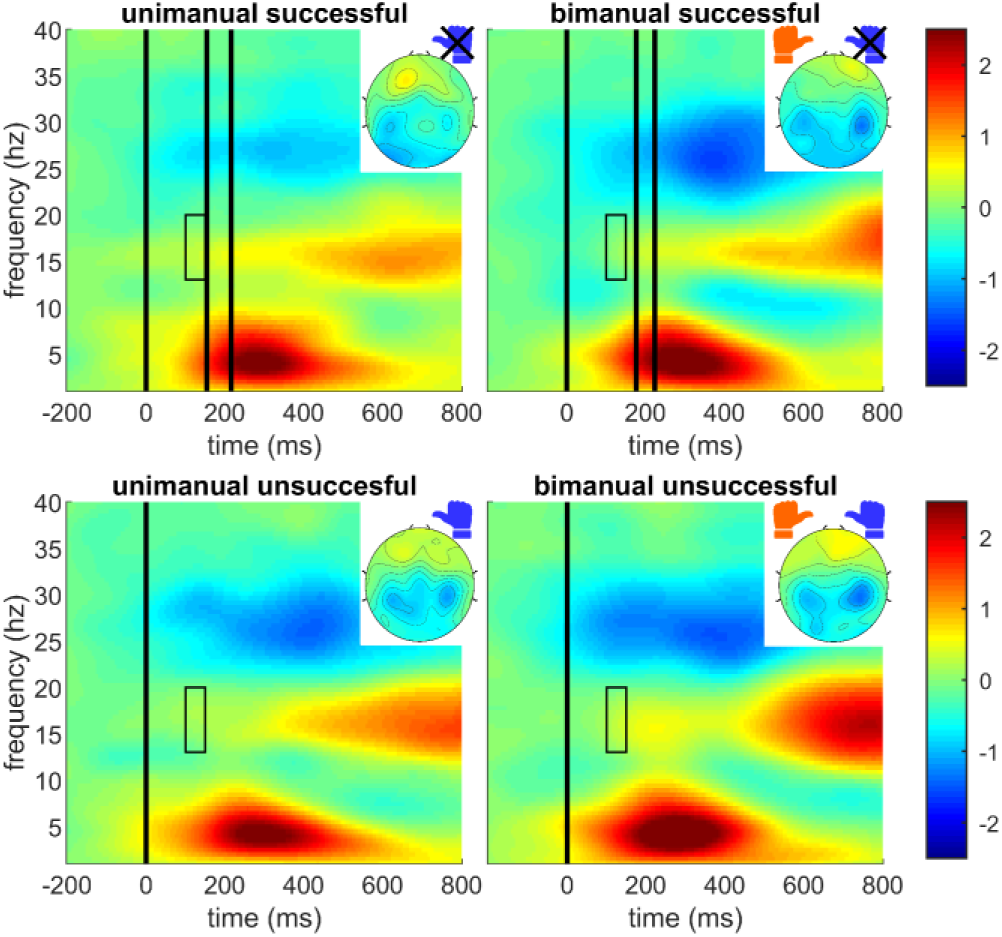
Stop trial time-frequency results time-locked to the stop signal. The images are of the average signal from frontocentral (F1, Fz, F2, FC1, FCz, FC2) electrodes. Vertical lines indicate, from left to right, stop signal onset, average prEMG latency, and average SSRT (only stop onset marked on unsuccessful stop trials). Rectangles indicate the time− and frequencies-of-interest included in the frontal beta analysis (13-20 Hz at 100-150 ms). The topographies show the distribution of beta values across all electrodes at the extracted time-window. The data for left and right hand trials are averaged in all panels, with the electrodes flipped in relevant conditions so that the depicted left hemisphere is contralateral to the stop hand. This is also reflected by the hand symbols, where blue refers to the stop-hand and orange refers to the response-hand, and the crosses above the blue hands indicate that these responses were successfully inhibited.

#### Summary I

We found no evidence for distinct global and selective stopping mechanisms. There were no differences between unimanual and bimanual stopping in behavior, corticomotor excitability for the stopped hand, or frontal beta power. The delayed prEMG peak in the bimanual task relative to the unimanual task, when considered on its own, is consistent with the dual-mechanism model. However, this delay was greatly diminished after accounting for differences in the prEMG onsets, a marker of the go process’s latency. The prevalent interference effect in the bimanual task and the suppression of the unselected hand during stopping, evident in reduced MEP amplitudes, support a single global inhibition mechanism model.

### 2) Does motor preparation influence subsequent inhibition success?

We hypothesized that motor cortical activity prior to the stop signal is associated with subsequent stopping success. Specifically, we expected weaker sensorimotor mu and beta desynchronization on successful than unsuccessful stop trials prior to stop signal presentation. Such a pattern could arise from a less mature go process and/or the influence of proactive control.

#### Weaker hand-specific mu desynchronization on bimanual successful stop trials

Mu desynchronization was prominent in all trials (Figure 6A; one-sample t-tests against zero: BF = 112.30; bimanual BF = 29.86). In the unimanual task, there was only a main effect of LOCATION (BF = 3.09*10^5^) indicating that mu desynchronization was stronger in the contralateral hemisphere, but no effect of SUCCESS (BF = 0.66). In the bimanual task, there was a weak interaction of SUCCESS and LOCATION (BF = 1.43). The post hoc tests indicated that mu desynchronization was weaker prior to successful stop trials in the hemisphere contralateral to the stopped hand, compared to the hemisphere contralateral to the response hand (Figure 6B, right panel; BF = 9.88). In contrast, mu activity was similar in both hemispheres on unsuccessful stop trials (BF = 0.25).

**Figure 6.**
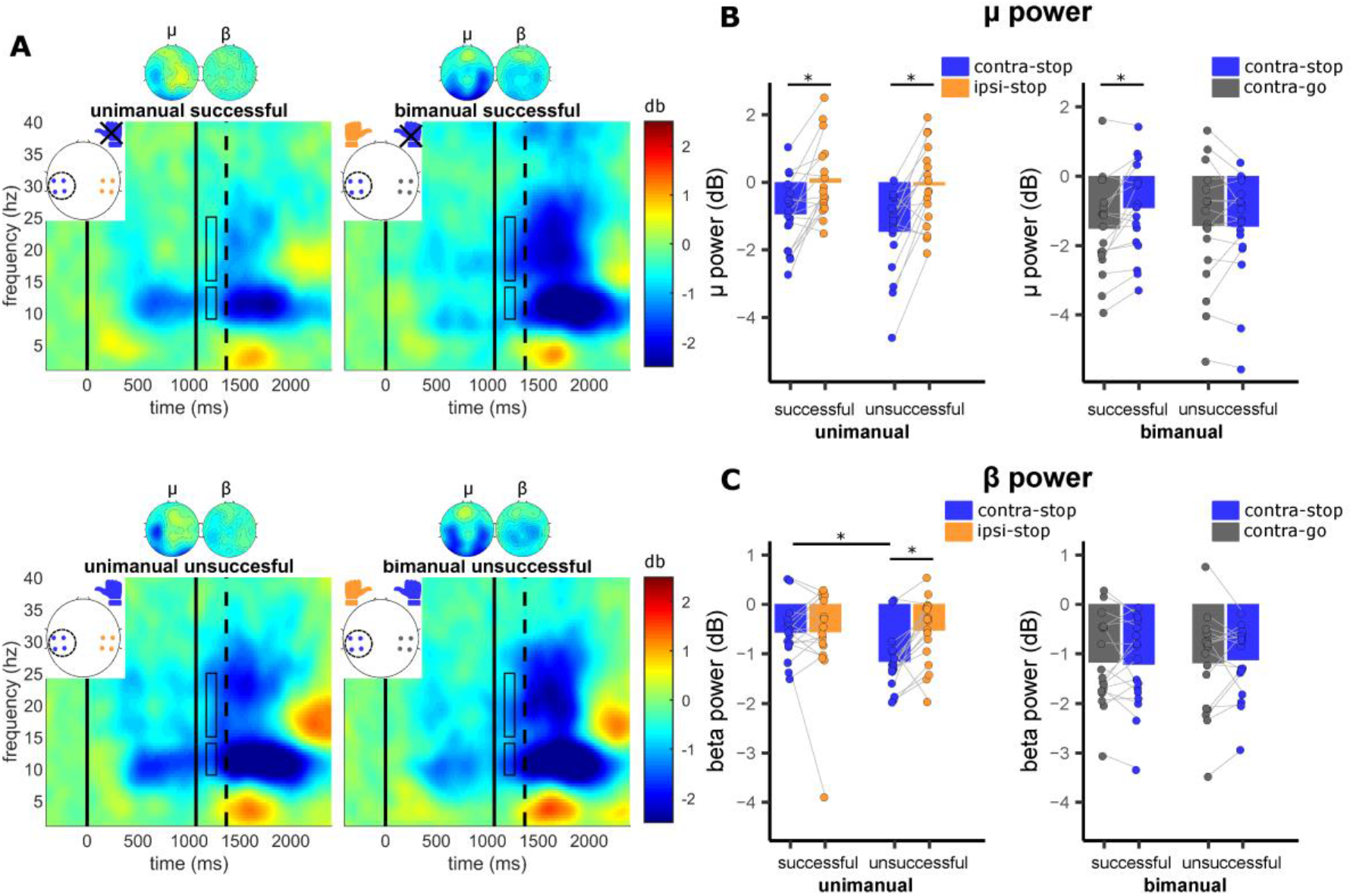
Time-frequency results on stop trials, time-locked to the cue onset. The spectrograms represent the average signal from the circled electrodes on the white topographical plots. The electrode colors correspond to the hemispheres shown in B and C; the blue hands symbolize the stop response (top: successful, and bottom: unsuccessful), and the orange hands symbolize the (correctly) executed response. The three black vertical lines represent, from left to right, cue onset, go onset, and average stop signal onset. Rectangles indicate the time- and frequencies-of-interest included in the analysis (9-14 Hz (mu) and 15-25 Hz (beta) at 100-200 ms after go onset). The topographies show the distribution of mu and beta values across all electrodes at the extracted time-window. **B**. Average mu power. **C**. Average beta power. * condition differences with BF > 3.

#### Weaker contralateral beta desynchronization on unimanual successful stop trials

The post-go beta power was also reduced in both tasks (Figure 6A; one sample t-tests against zero: unimanual BF = 1.72*10^4^; bimanual BF = 1206.55), yet the effects observed here differed from those observed in the mu band. Relevant effects were limited to the unimanual task where there was an interaction of SUCCESS and LOCATION (BF = 2.86; Figure 6C). The latter reflected the finding that beta desynchronization in the contralateral hemisphere was weaker prior to successful stop trials compared to unsuccessful stop trials (BF = 27.88). Further, beta desynchronization was stronger in contra-than ipsilateral hemisphere prior to unsuccessful (BF = 78.12), but not prior to successful stop trials (BF = 0.25).

#### Summary II

The state of motor activity prior to stop signal onset influenced subsequent stopping success. In the unimanual task, beta desynchronization was weaker prior to successful compared to unsuccessful stopping. Selective stopping in the bimanual task exhibited a distinct pattern of mu lateralization: mu desynchronization contralateral to the stopped hand was weaker than in the hemisphere contralateral to the responding hand prior to successful stop trials, while no lateralization effect was observed prior to the unsuccessful stop trials.

### 3) Is motor preparation affected by proactive control?

The effect of proactive control on motor activity was tested by contrasting the trials with cues indicating that the prepared response may need to be stopped (MS) to the trials in which the cue indicated the certain execution of the prepared response (CG). We expected the MS cue to lead to slower go RTs, altered MEP amplitudes during the cue-delay period, and reduced motor mu and beta desynchronization.

#### Delayed go RTs in the MS cue condition

Relative to the CG condition, RTs were delayed by ~150 ms in the MS condition (Table 3; BF = 4.28*10^10^). However, there was no effect of the TASK on go RTs (BF = 0.35), nor an interaction of TASK and CUE (BF = 0.28). Importantly, there were no systematic RT differences in the bimanual go trials of the MS condition between the hand that was cued to respond compared to the hand that was cued to potentially stop (both 532 ms; BF = 0.24). In sum, proactive control led to substantial response slowing of comparable magnitude in both tasks.

#### Hand-specific cue-related modulation of the MEPs in the bimanual task, but no cueing effects in the unimanual task

Turning to the TMS data, we observed neither suppression nor excitation of corticospinal excitability compared to the baseline during the delay period for either type of cue (all BFs <= 0.78; Figure 4D).

In the unimanual task, MEPs elicited in the unselected hand were smaller than those elicited in the selected hand (HAND: BF = 5.54), with no difference between the CG and MS conditions (CUE: BF = 0.25). In the bimanual task, a one-way rmANOVA comparing the three conditions, CG, MS go hand, MS maybe stop hand, showed weak evidence for the main effect (BF = 1.44). Post hoc tests indicated that MEP amplitudes were reduced in the hand that was cued for stopping in the MS condition compared to the CG condition (BF = 4.77) and compared to when the hand was not cued for stopping in the MS condition (BF = 1.20). There were no differences between the responding hands in the CG and MS condition (BF = 0.25).

In sum, there were no differences between the CG and MS responding hand in the unimanual task. However, the MEPs in the hand that was cued to stop were reduced compared to the responding hand in both cue conditions.

#### Reduced mu and beta desynchronization in the MS cue condition

Sensorimotor mu and beta desynchronization were tested between the cue types and hemispheres. To evaluate the time course of the desynchronization effects, we included time-window (pre- vs post-go signal) as an additional factor in the analyses (Figure 7A). One-sample t-tests against zero confirmed the presence of mu desynchronization both in the pre-go (unimanual BF = 739.34; bimanual BF = 144.97) and post-go period (unimanual BF = 290.63; bimanual BF = 138.89).

**Figure 7.**
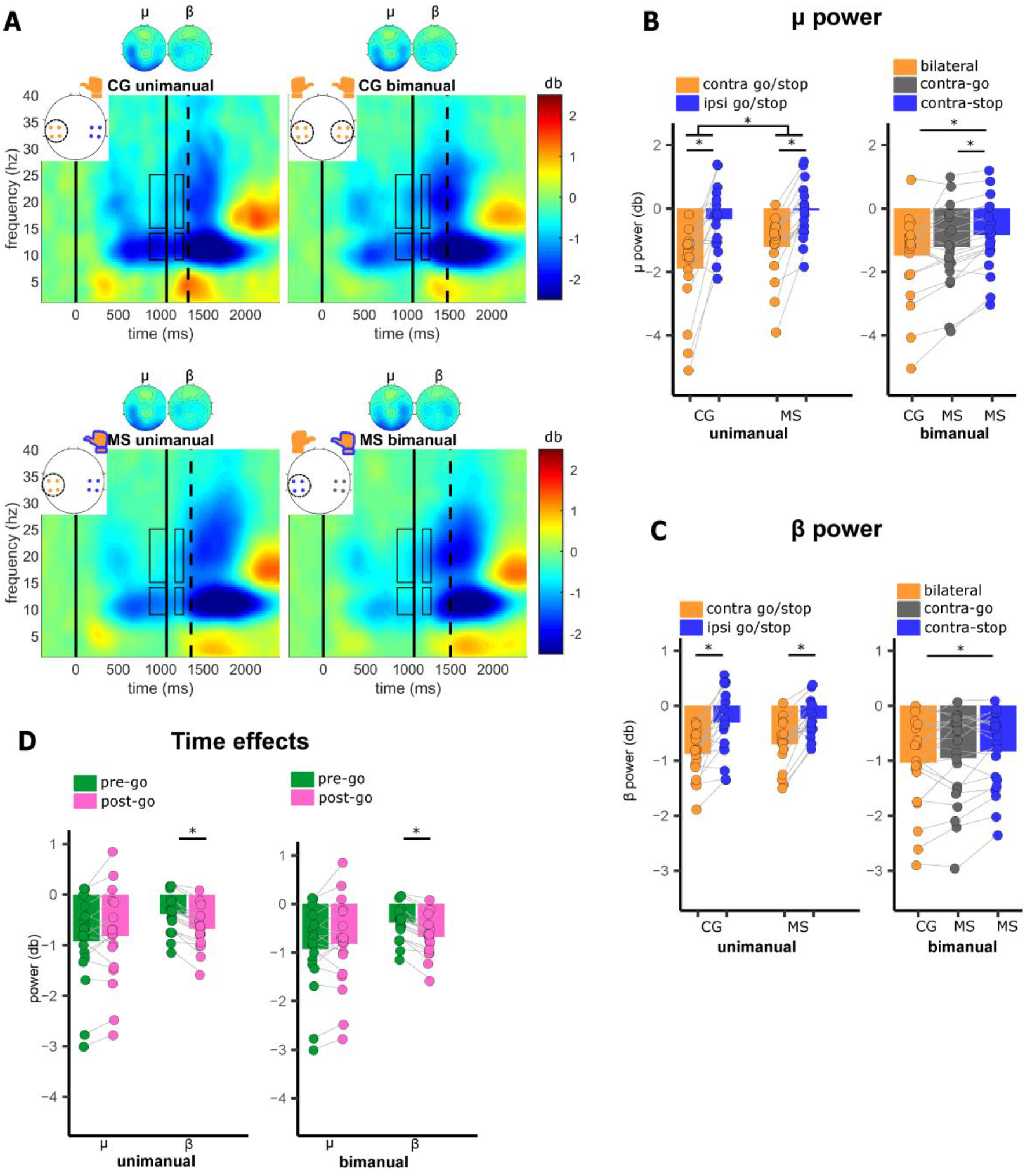
**A**. Go trial time-frequency results, time-locked to the cue onset. The spectrograms depict the averaged signal of the encircled electrodes on the white topographical plots. The electrode colors correspond to the hemispheres shown in B and C; all hand symbols are orange depicting that the response was always given, but the blue outline indicates which hands were additionally prepared for the maybe stop response. Vertical lines indicate, from left to right: cue onset, go onset, and average go-EMG onset. Rectangles indicate the time- and frequencies-of-interest included in the analysis (9-14 Hz (mu) and 15-25 Hz (beta) at −200-0 and 100-200 ms relative to go onset. The topographies show the distribution of mu and beta values across all electrodes averaged over both pre-go and post-go time-windows. **B**. Average mu power (averaged over pre-go and post-go period). **C**. Average beta power (averaged over pre-go and post-go period). **D**. Pre-go and post-go mu and beta power, averaged over hemispheres and cue conditions. CG – certain go, MS – maybe stop. *represent condition differences with BF > 3.

In the unimanual task, mu desynchronization was stronger following CG cues compared to MS cues (CUE: BF = 6.45), and stronger over the contralateral hemisphere compared to the ipsilateral hemisphere (LOCATION: BF = 7.69*10^15^; Figure 7B, left panel). This desynchronization was evident during the delay period and persisted following the imperative stimulus with no difference between the pre- and post-go periods (Figure 7D; TIME: BF = 0.21).

In the bimanual task, since both hands are responding simultaneously, the terms ipsilateral and contralateral are ambiguous. Therefore, both cue and hemispheric effects were tested with a rmANOVA with a combined CUE-LOCATION factor (levels CG bilateral, MS contra-go, MS contra-stop). This analysis revealed a significant main effect (BF = 2.94*10^4^; Figure 7B, right panel). The post-hoc tests indicated that the mu desynchronization was stronger in the CG than in the MS contra-stop condition (BF = 10.82) and in MS contra-go than in MS contra-stop hemisphere (BF = 174.45), paralleling the changes in the pre-go MEP amplitudes. There was no main effect of TIME (BF = 0.78, Figure 7D).

#### Hand-specific beta modulation in the bimanual task, but no cue effects in the unimanual task

Beta desynchronization was also evident in both tasks (Figure 7A and 7C) during pre-go (one-sample t-tests against zero: unimanual BF = 2952.96; bimanual BF = 273.97) and post-go periods (unimanual BF = 1.47*10^5^; bimanual BF = 2025.58). Beta desynchronization was stronger later in the trial, after the go signal (Figure 7D; TIME unimanual: BF = 205.36; bimanual: BF = 9.36*10^7^). Beta desynchronization was stronger in the contra-than in the ipsilateral hemisphere, relative to the selected hand in the unimanual task (LOCATION: BF = 2.79*10^10^). There were no cue-effects in the unimanual task (CUE: BF = 0.59), but there was minor evidence for cue effects in the bimanual task (CUE-LOCATION: BF = 1.50). The post-hoc tests revealed that the beta desynchronization contralateral to the stopping hand in the MS cue condition was weaker in the MS cue compared to the CG cue (BF = 4.80).

#### Summary III

Proactive control influenced motor preparation, evident in pronounced RT increases on MS trials in both tasks. MEPs during the delay period were attenuated in the MS condition in the bimanual task, but only in the hand that was cued to maybe-stop. In the unimanual condition, MEP amplitudes in the pre-go period did not differ between the two types of cues. Thus, the MEP changes were restricted to the condition in which participants prepared for selective stopping, consistent with previous studies (Claffey et al., 2010; Cai et al., 2011). As expected, mu desynchronization was weaker in MS than in CG trials, paralleling the RT differences. Moreover, mu desynchronization was weaker in the hemisphere corresponding to the hand cued for possible stopping compared to the freely responding hand in the bimanual task. This laterality effect is consistent with the observed pattern of task-specific MEP modulation.

## Discussion

### Single inhibition mechanism with predominantly global extent

We evaluated the explanatory power of two models of response inhibition. As summarized in Table 5, the results were more consistent with the single global inhibition model as indicated by stopping interference in the behavior and EMG time-courses, and the suppressed MEPs of the unselected hand after the stop signal. There were no differences in the SSRTs between uni- and bimanual stopping, stopped hand MEP amplitudes, or frontal beta power. These results are at odds with the predictions of the dual-mechanism model in which inhibition can be selectively directed. An initial difference in prEMG between the tasks diminished after controlling for dependencies between go and stop processes. As the single trial analysis further indicated dependencies between the prEMG onset time and stopping, we speculate that the hypothesized delays previously associated with selective inhibition may be driven by differences in the go process or interactions between going and stopping.

**Table 5.** Predictions regarding putative inhibition indices (SSRT, prEMG, MEP amplitudes, frontal beta power) in the context of the two models. The marks on the right indicate whether these predictions were confirmed (✓) or not confirmed (✗) by the data. The ✗/✓ marks that the results were inconclusive.

|  |  |
| --- | --- |
| **Global inhibition** |  |
| Interference in the responding hand RT in bimanual successful stop trials | ✓ |
| Interference in the responding hand EMG in the bimanual successful stop trials | ✓ |
| Suppressed unselected hand post-stop MEP amplitude in the unimanual task | ✓ |
| **Dual-mechanism** |  |
| Delayed SSRT in the bimanual task | ✗ |
| Delayed prEMG peak latency in the bimanual task | ✗/✓ |
| Weaker post-stop MEP suppression in the bimanual task | ✗ |
| Reduced post-stop beta power in the bimanual task | ✗ |

While the preponderance of evidence supports a single global inhibition mechanism, a few effects were indicative of selective stopping. We devised a novel single-trial marker, the Δpeak, for quantifying interference in the bimanual task. Despite the prevalent interference in the EMG at the group level, the single-trial Δpeak indicated large variability, including trials with no interference. This is in line with previous findings showing that interference can be eliminated under specific conditions (Xu et al., 2015) or when a small delay is required between the bimanual left and right hand responses (in contrast to simultaneous ones), which necessitates a decoupling of the bimanual response (Wadsley et al., 2019). These results highlight that the stop-restart behavior in the bimanual stop signal task is not mandatory, but a preferred option when selectivity is not incentivized by the task requirements. Our results further indicate that stopping consists of a mix of processes, including trials where the entire action plan was aborted (large Δpeak), trials where there was selective response preparation and/or inhibition (small or no Δpeak), and trials where the motor plan was aborted before motor initiation (no prEMG; De Jong et al., 1990; McGarry and Franks, 1997).

These results motivate an alternative to the global inhibition model, one that highlights flexible control, the extent and speed of which vary on a trial-by-trial basis depending on the motivational and task context. Inhibition may be one component of a continually revised motor plan, incorporating simultaneous facilitatory and inhibitory drives. These dynamics could depend on the wider sensorimotor control system spanning the dorsal attentional, motor, and premotor areas (Cisek and Kalaska, 2010; Mirabella, 2014). One overarching function of cortical inhibition in this model could be to set the gain for motor representations, with the extent of inhibition changing according to uncertainty about the response alternatives or requirements for proactive control (Greenhouse et al., 2015).

### Proactive control, motor preparation, and response inhibition

Proactive control biases sensorimotor and attentional systems in expectation of a stop signal (Elchlepp et al., 2016; Langford et al., 2016). Our results speak to the role of anticipatory regulation on the motor system. First, we observed that motor cortical activity differed between successful and unsuccessful stopping prior to stop signals. Second, we found smaller MEPs in the hand that was cued to potentially stop, which is in line with previous findings of hand-specific modulation of anticipatory corticomotor excitability (Claffey et al., 2010; Cai et al., 2011) or short-interval intracortical inhibition (Cirillo et al., 2018). Further, sensory-motor EEG activity changed with proactive cues in a hand-specific way, paralleling the pre-go MEP modulation in the bimanual task. These results suggest functional interactions between proactive control, motor preparation, and response inhibition. Successful inhibition may arise reflexively with respect to acquired stimulus-stop associations with sensory, attentional, and motor parameters pre-set by proactive control (Verbruggen et al., 2014a, 2014b).

### EEG signatures of response inhibition and motor preparation

A number of EEG markers have been proposed to index response inhibition. However, most of these occur relatively late in the trial. The prEMG latencies in this study are consistent with recent findings showing that inhibition occurs around 140-180 ms after the stop signal (Raud and Huster, 2017; Atsma et al., 2018; Hannah et al., 2019; Huster et al., 2019; Jana et al., 2020; Raud et al., 2020; Thunberg et al., 2020). Beta power increases in the right inferior frontal gyrus and pre-supplementary motor area within 100 ms following the stop signal (Swann et al., 2009, 2012). However, we did not find increased beta in the time-window of 100-150 ms after stopping. Moreover, beta activity did not differentiate between selective and global stopping, nor between successful and unsuccessful stop trials. The latter findings are in line with intracranial recordings of beta power in the pre-SMA (Swann et al., 2012) and a study showing that the number of frontal beta bursts (a potentially more sensitive marker than power fluctuations) did not differ between successful and unsuccessful stopping before 200 ms (Wessel, 2020). As such, a fast, reactive inhibition mechanism may invariably be engaged whenever a stop signal is detected, while the success of stopping may depend on the perceptual speed and the contemporaneous state of the motor cortex.

Regarding the cortical activity within the sensorimotor system, we observed weaker mu desynchronization accompanied by delayed go RTs in the cued condition. In the bimanual task, we observed weaker mu and beta desynchronization in the hemisphere contralateral to the maybe-stop hand, paralleled by reduced MEP amplitudes in the hand cued for stopping. Weaker desynchronization may therefore reflect the engagement of proactive control or its downstream effects on the motor system, in line with the hypothesis that cortical oscillations in these frequency bands reflect more general inhibitory functions, not only those limited to the motor cortex (Jensen and Mazaheri, 2010). Accordingly, Muralidharan et al. (2019) reported a power increase relative to baseline in the hemisphere contralateral to the stop hand in a frequency range incorporating both mu and beta bands, which they suggest was reflective of proactive selective inhibition of the motor cortex. Here, mu and beta were negative relative to the baseline in both hemispheres. It is thus important to acknowledge the possibility of relative differences between the two hemispheres arising due to the lateralized disinhibition of the portion of the upcoming movement that can be freely executed.

## Limitations

While our results agree with the previous reports on uni- and bimanual stopping, there are important differences to note. The validity of the SSRT relies on the assumption of independent go and stop processes, yet we demonstrated and quantified violations of this assumption at the single trial level. The validity of existing methods for SSRT calculation has been called into question based on observations of similar violations (Ozyurt et al., 2003; Bissett and Logan, 2014; Gulberti et al., 2014; Verbruggen and Logan, 2015), biases produced by subtle changes in the go RT distribution (Verbruggen et al., 2013), and the inability to account for attentional lapses (Matzke et al., 2017; Heathcote et al., 2019). Altogether, these findings call for a re-conceptualization of SSRT differences observed between different task conditions.

MEPs were measured from a single effector at two time-points: before the go and after the stop stimulus. This is suboptimal both for measuring the spread of inhibition to task unrelated effectors and for sampling the temporal evolution of corticomotor excitability. For example, even though we observed reduced post-stop MEPs in the unselected hand as a putative indicator of global inhibition, interhemispheric inhibition during response selection could also influence MEPs. Further, we did not replicate the robust preparatory inhibition of corticomotor excitability in a hand selected for a forthcoming response reported in a number of studies (e.g., Duque et al., 2017). The same group has recently shown that preparatory inhibition does not generalize across different task contexts (Quoilin et al., 2019), and several design features of our study differed from previous studies (e.g., the inclusion of stop trials, the target muscle, stimulation hemisphere, pulse timing, and inter-stimulation interval among others). Altogether, separating the various forms of inhibitory control during response generation and inhibition, as well as their effects on selected, unselected, and task-irrelevant effectors offers a rich ground for future research.

## Conclusions

Our results favored a single inhibition mechanism over a model involving independent selective and global mechanisms. We further observed that the success of stopping differed according to the state of sensorimotor activity preceding the stop signal. Moreover, the scalp distribution of sensorimotor mu and beta activity corresponded to TMS measurements of hand-specific reduction of corticomotor excitability in the hand cued for possible stopping. The results, taken as a whole, are consistent with a model emphasizing the global extent of inhibition. However, the novel use of EMG activity to quantify stopping interference at the single trial level demonstrated that selective stopping can be achieved in certain trials, pointing to greater flexibility of inhibitory mechanism in terms of their spatial extent.

## Acknowledgments

This work was supported by the University of Oslo and the Peder Sather Center, an International Research and Educational Collaboration between UC Berkeley and Norway. RBI was supported by a grant from the National Institutes of Health, NS092079. IG was supported by a grant from the National Institutes of Health, TR002370. We thank Claudia Tischler and Vincent Ngo for their assistance during data collection, and Assaf Breska and Nicole Swann for their helpful comments.

## References

Aron AR (2011) From reactive to proactive and selective control: developing a richer model for stopping inappropriate responses. Biol Psychiatry 69:e55–68.

Aron AR, Verbruggen F (2008) Stop the presses: dissociating a selective from a global mechanism for stopping. Psychol Sci 19:1146–1153.

Atsma J, Maij F, Gu C, Medendorp WP, Corneil BD (2018) Active Braking of Whole-Arm Reaching Movements Provides Single-Trial Neuromuscular Measures of Movement Cancellation. J Neurosci Off J Soc Neurosci 38:4367–4382.

Badry R, Mima T, Aso T, Nakatsuka M, Abe M, Fathi D, Foly N, Nagiub H, Nagamine T, Fukuyama H (2009) Suppression of human cortico-motoneuronal excitability during the Stop-signal task. Clin Neurophysiol Off J Int Fed Clin Neurophysiol 120:1717–1723.

Band GPH, van der Molen MW, Logan GD (2003) Horse-race model simulations of the stop-signal procedure. Acta Psychol (Amst) 112:105–142.

Bestmann S, Krakauer JW (2015) The uses and interpretations of the motor-evoked potential for understanding behaviour. Exp Brain Res 233:679–689.

Bissett PG, Logan GD (2014) Selective stopping? Maybe not. J Exp Psychol Gen 143:455–472.

Boucher L, Palmeri TJ, Logan GD, Schall JD (2007) Inhibitory control in mind and brain: an interactive race model of countermanding saccades. Psychol Rev 114:376–397.

Brinkman L, Stolk A, Marshall TR, Esterer S, Sharp P, Dijkerman HC, de Lange FP, Toni I (2016) Independent Causal Contributions of Alpha− and Beta-Band Oscillations during Movement Selection. J Neurosci Off J Soc Neurosci 36:8726–8733.

Cai W, Oldenkamp CL, Aron AR (2011) A proactive mechanism for selective suppression of response tendencies. J Neurosci Off J Soc Neurosci 31:5965–5969.

Cai W, Oldenkamp CL, Aron AR (2012) Stopping speech suppresses the task-irrelevant hand. Brain Lang 120:412–415.

Chaumon M, Bishop DVM, Busch NA (2015) A practical guide to the selection of independent components of the electroencephalogram for artifact correction. J Neurosci Methods 250:47–63.

Chikazoe J, Jimura K, Hirose S, Yamashita K, Miyashita Y, Konishi S (2009) Preparation to inhibit a response complements response inhibition during performance of a stop-signal task. J Neurosci Off J Soc Neurosci 29:15870–15877.

Cirillo J, Cowie MJ, MacDonald HJ, Byblow WD (2018) Response inhibition activates distinct motor cortical inhibitory processes. J Neurophysiol 119:877–886.

Cisek P, Kalaska JF (2010) Neural mechanisms for interacting with a world full of action choices. Annu Rev Neurosci 33:269–298.

Claffey MP, Sheldon S, Stinear CM, Verbruggen F, Aron AR (2010) Having a goal to stop action is associated with advance control of specific motor representations. Neuropsychologia 48:541–548.

Cowie MJ, MacDonald HJ, Cirillo J, Byblow WD (2016) Proactive modulation of long-interval intracortical inhibition during response inhibition. J Neurophysiol 116:859–867.

Coxon JP, Stinear CM, Byblow WD (2007) Selective inhibition of movement. J Neurophysiol 97:2480–2489.

De Jong R, Coles MG, Logan GD, Gratton G (1990) In search of the point of no return: the control of response processes. J Exp Psychol Hum Percept Perform 16:164–182.

Delorme A, Makeig S (2004) EEGLAB: an open source toolbox for analysis of single-trial EEG dynamics including independent component analysis. J Neurosci Methods 134:9–21.

Duque J, Greenhouse I, Labruna L, Ivry RB (2017) Physiological Markers of Motor Inhibition during Human Behavior. Trends Neurosci 40:219–236.

Elchlepp H, Lavric A, Chambers CD, Verbruggen F (2016) Proactive inhibitory control: A general biasing account. Cognit Psychol 86:27–61.

Greenhouse I, Oldenkamp CL, Aron AR (2012) Stopping a response has global or nonglobal effects on the motor system depending on preparation. J Neurophysiol 107:384–392.

Greenhouse I, Sias A, Labruna L, Ivry RB (2015) Nonspecific Inhibition of the Motor System during Response Preparation. J Neurosci Off J Soc Neurosci 35:10675–10684.

Gulberti A, Arndt PA, Colonius H (2014) Stopping eyes and hands: evidence for non-independence of stop and go processes and for a separation of central and peripheral inhibition. Front Hum Neurosci 8:61.

Hannah R, Muralidharan V, Sundby KK, Aron AR (2019) Temporally-precise disruption of prefrontal cortex informed by the timing of beta bursts impairs human action-stopping. bioRxiv:843557.

Heathcote A, Lin Y-S, Reynolds A, Strickland L, Gretton M, Matzke D (2019) Dynamic models of choice. Behav Res Methods 51:961–985.

Huster RJ, Messel MS, Thunberg C, Raud L (2019) The P300 as marker of inhibitory control –fact or fiction? bioRxiv:694216.

Jahfari S, Stinear CM, Claffey M, Verbruggen F, Aron AR (2010) Responding with restraint: what are the neurocognitive mechanisms? J Cogn Neurosci 22:1479–1492.

Jahfari S, Verbruggen F, Frank MJ, Waldorp LJ, Colzato L, Ridderinkhof KR, Forstmann BU (2012) How preparation changes the need for top-down control of the basal ganglia when inhibiting premature actions. J Neurosci Off J Soc Neurosci 32:10870–10878.

Jana S, Hannah R, Muralidharan V, Aron AR (2020) Temporal cascade of frontal, motor and muscle processes underlying human action-stopping van den Wildenberg W, Ivry RB, van den Wildenberg W, Huster R, Bissett PG, eds. eLife 9:e50371.

Jensen O, Mazaheri A (2010) Shaping functional architecture by oscillatory alpha activity: gating by inhibition. Front Hum Neurosci 4:186.

Kayser J, Tenke CE (2006) Principal components analysis of Laplacian waveforms as a generic method for identifying ERP generator patterns: I. Evaluation with auditory oddball tasks. Clin Neurophysiol Off J Int Fed Clin Neurophysiol 117:348–368.

Krämer UM, Knight RT, Münte TF (2011) Electrophysiological evidence for different inhibitory mechanisms when stopping or changing a planned response. J Cogn Neurosci 23:2481–2493.

Langford ZD, Krebs RM, Talsma D, Woldorff MG, Boehler CN (2016) Strategic down-regulation of attentional resources as a mechanism of proactive response inhibition. Eur J Neurosci.

Lavallee CF, Meemken MT, Herrmann CS, Huster RJ (2014) When holding your horses meets the deer in the headlights: time-frequency characteristics of global and selective stopping under conditions of proactive and reactive control. Front Hum Neurosci 8:994.

Liebrand M, Kristek J, Tzvi E, Krämer UM (2018) Ready for change: Oscillatory mechanisms of proactive motor control. PloS One 13:e0196855.

Liebrand M, Pein I, Tzvi E, Krämer UM (2017) Temporal Dynamics of Proactive and Reactive Motor Inhibition. Front Hum Neurosci 11:204.

Logan GD, Cowan WB (1984) On the ability to inhibit thought and action: A theory of an act of control. Psychol Rev 91:295–327.

Macdonald HJ, Coxon JP, Stinear CM, Byblow WD (2014) The Fall and Rise of Corticomotor Excitability with Cancellation and Reinitiation of Prepared Action. J Neurophysiol.

Macdonald HJ, Stinear CM, Byblow WD (2012) Uncoupling response inhibition. J Neurophysiol 108:1492–1500.

Majid DSA, Cai W, George JS, Verbruggen F, Aron AR (2012) Transcranial magnetic stimulation reveals dissociable mechanisms for global versus selective corticomotor suppression underlying the stopping of action. Cereb Cortex N Y N 1991 22:363–371.

Matzke D, Love J, Heathcote A (2017) A Bayesian approach for estimating the probability of trigger failures in the stop-signal paradigm. Behav Res Methods 49:267–281.

Mazaheri A, Nieuwenhuis ILC, van Dijk H, Jensen O (2009) Prestimulus alpha and mu activity predicts failure to inhibit motor responses. Hum Brain Mapp 30:1791–1800.

McGarry T, Franks IM (1997) A horse race between independent processes: evidence for a phantom point of no return in preparation of a speeded motor response. J Exp Psychol Hum Percept Perform 23:1533–1542.

Mirabella G (2014) Should I stay or should I go? Conceptual underpinnings of goal-directed actions. Front Syst Neurosci 8:206.

Munakata Y, Herd SA, Chatham CH, Depue BE, Banich MT, O’Reilly RC (2011) A unified framework for inhibitory control. Trends Cogn Sci 15:453–459.

Muralidharan V, Yu X, Cohen MX, Aron AR (2019) Preparing to Stop Action Increases Beta Band Power in Contralateral Sensorimotor Cortex. J Cogn Neurosci 31:657–668.

Neuper C, Wörtz M, Pfurtscheller G (2006) ERD/ERS patterns reflecting sensorimotor activation and deactivation. Prog Brain Res 159:211–222.

Oldfield RC (1971) The assessment and analysis of handedness: the Edinburgh inventory. Neuropsychologia 9:97–113.

Ozyurt J, Colonius H, Arndt PA (2003) Countermanding saccades: evidence against independent processing of go and stop signals. Percept Psychophys 65:420–428.

Picazio S, Veniero D, Ponzo V, Caltagirone C, Gross J, Thut G, Koch G (2014) Prefrontal control over motor cortex cycles at beta frequency during movement inhibition. Curr Biol CB 24:2940–2945.

Quoilin C, Fievez F, Duque J (2019) Preparatory inhibition: Impact of choice in reaction time tasks. Neuropsychologia 129:212–222.

Raud L, Huster RJ (2017) The Temporal Dynamics of Response Inhibition and their Modulation by Cognitive Control. Brain Topogr 30:486–501.

Raud L, Westerhausen R, Dooley N, Huster RJ (2020) Differences in unity: The go/no-go and stop signal tasks rely on different mechanisms. NeuroImage 210:116582.

Ruddy KL, Woolley DG, Mantini D, Balsters JH, Enz N, Wenderoth N (2018) Improving the quality of combined EEG-TMS neural recordings: Introducing the coil spacer. J Neurosci Methods 294:34–39.

Schall JD, Palmeri TJ, Logan GD (2017) Models of inhibitory control. Philos Trans R Soc Lond B Biol Sci 372.

Smittenaar P, Guitart-Masip M, Lutti A, Dolan RJ (2013) Preparing for selective inhibition within frontostriatal loops. J Neurosci Off J Soc Neurosci 33:18087–18097.

Smittenaar P, Rutledge RB, Zeidman P, Adams RA, Brown H, Lewis G, Dolan RJ (2015) Proactive and Reactive Response Inhibition across the Lifespan. PloS One 10:e0140383.

Stuphorn V, Emeric EE (2012) Proactive and reactive control by the medial frontal cortex. Front Neuroengineering 5:9.

Swann N, Tandon N, Canolty R, Ellmore TM, McEvoy LK, Dreyer S, DiSano M, Aron AR (2009) Intracranial EEG reveals a time− and frequency-specific role for the right inferior frontal gyrus and primary motor cortex in stopping initiated responses. J Neurosci Off J Soc Neurosci 29:12675–12685.

Swann NC, Cai W, Conner CR, Pieters TA, Claffey MP, George JS, Aron AR, Tandon N (2012) Roles for the pre-supplementary motor area and the right inferior frontal gyrus in stopping action: electrophysiological responses and functional and structural connectivity. NeuroImage 59:2860–2870.

Thunberg C, Messel MS, Raud L, Huster RJ (2020) tDCS over the inferior frontal gyri and visual cortices did not improve response inhibition. Sci Rep 10:7749.

van den Wildenberg WPM, Burle B, Vidal F, van der Molen MW, Ridderinkhof KR, Hasbroucq T (2010) Mechanisms and dynamics of cortical motor inhibition in the stop-signal paradigm: a TMS study. J Cogn Neurosci 22:225–239.

Verbruggen F et al. (2019) A consensus guide to capturing the ability to inhibit actions and impulsive behaviors in the stop-signal task Frank MJ, Badre D, Egner T, Swick D, eds. eLife 8:e46323.

Verbruggen F, Best M, Bowditch WA, Stevens T, McLaren IPL (2014a) The inhibitory control reflex. Neuropsychologia 65:263–278.

Verbruggen F, Chambers CD, Logan GD (2013) Fictitious inhibitory differences: how skewness and slowing distort the estimation of stopping latencies. Psychol Sci 24:352–362.

Verbruggen F, Logan GD (2015) Evidence for capacity sharing when stopping. Cognition 142:81–95.

Verbruggen F, McLaren IPL, Chambers CD (2014b) Banishing the Control Homunculi in Studies of Action Control and Behavior Change. Perspect Psychol Sci J Assoc Psychol Sci 9:497–524.

Wadsley CG, Cirillo J, Byblow WD (2019) Between-hand coupling during response inhibition. J Neurophysiol.

Wagner J, Wessel JR, Ghahremani A, Aron AR (2018) Establishing a Right Frontal Beta Signature for Stopping Action in Scalp EEG: Implications for Testing Inhibitory Control in Other Task Contexts. J Cogn Neurosci 30:107–118.

Wessel JR (2020) β-Bursts Reveal the Trial-to-Trial Dynamics of Movement Initiation and Cancellation. J Neurosci 40:411–423.

Wessel JR, Aron AR (2017) On the Globality of Motor Suppression: Unexpected Events and Their Influence on Behavior and Cognition. Neuron 93:259–280.

Wessel JR, Reynoso HS, Aron AR (2013) Saccade suppression exerts global effects on the motor system. J Neurophysiol 110:883–890.

Xu J, Westrick Z, Ivry RB (2015) Selective inhibition of a multicomponent response can be achieved without cost. J Neurophysiol 113:455–465.

